# Oxidative Stress and Tissue Remodeling in a Rat Abdominal Seroma Model

**DOI:** 10.64898/2026.10.06.757126

**Authors:** Xiaochao Xia, Ann M. Nour, Kevin He, Adam Boukind, Yuhan Guo, Sida Hao, Batool Arif, Lauren Schellhardt, Mohamed A. Zayed, Andrew E. Gelman, Matthew D. Wood, Justin M. Sacks, Xiaowei Li

**Affiliations:** Division of Plastic and Reconstructive Surgery, Department of Surgery, Washington University School of Medicine, St. Louis, MO 63110; Department of Biomedical Engineering, McKelvey School of Engineering, Washington University in St. Louis, St. Louis, MO 63130; Department of Urology, Zhejiang Hospital of Integrated Traditional Chinese and Western Medicine, Hangzhou, Zhejiang, China; Section of Vascular Surgery, Department of Surgery, Washington University School of Medicine, St. Louis, MO 63110; Division of Molecular Cell Biology, Washington University School of Medicine, St. Louis, MO 63110; St. Louis Veterans Affairs Health Care System, St. Louis, MO 63106; Division of Cardiothoracic Surgery, Department of Surgery, Washington University School of Medicine, St. Louis, MO 63110

**Author notes:** Corresponding Authors: Justin M. Sacks, MD, Chief, Division of Plastic and Reconstructive Surgery Professor of Surgery, Sidney M. Jr. and Robert H. Shoenberg Chair, Washington University School of Medicine, St. Louis, MO 63110, Xiaowei Li, PhD, Assistant Professor of Surgery, Division of Plastic and Reconstructive Surgery Washington University School of Medicine, St. Louis, MO 63110.

**Keywords:** Seroma, Abdominoplasty, Oxidative stress, Stromal remodeling, Inflammation, Fibrosis

## Abstract

Seroma is a common complication following abdominoplasty and other procedures involving extensive soft tissue dissection. Although animal seroma models have been reported, the stromal pathology underlying post-abdominoplasty seroma, particularly the role of oxidative stress, remains poorly characterized. Here, we developed a rat abdominal seroma model by surgically separating the subcutaneous tissue from the underlying muscle fascia and investigated the temporal evolution of the associated stromal response. Seroma developed in 9 of 20 operated rats, corresponding to a formation rate of 45.0%. Longitudinal ultrasound demonstrated that fluid accumulation peaked around postoperative day (POD) 7 and largely resolved by POD30. Despite macroscopic fluid resolution, substantial stromal remodeling persisted. Compared with normal abdominal tissue, seroma-associated tissues at POD14 and POD35 exhibited increased collagen deposition, macrophage accumulation, oxidative injury, and vascular remodeling. Additionally, at POD35, macrophage accumulation and 4-hydroxynonenal staining were greater than at POD14, whereas collagen deposition and vascular density remained elevated with no significant differences between postoperative time points. To our knowledge, this study presents the first rat model specifically designed to reproduce post-abdominoplasty seroma and the first temporal characterization of oxidative stress within seroma-associated tissue. These findings reveal that seroma fluid resolution does not coincide with stromal recovery and identify persistent oxidative stress, inflammation, fibrosis, and vascular remodeling as important features of seroma pathology.

## Introduction

Seroma is the accumulation of serous fluid within a surgically created or traumatic tissue space and is a common complication following extensive soft tissue dissection^1^. It can delay wound healing, increase the risk of infection and wound dehiscence, and require repeated aspiration or drainage^1,2^. Clinically, seroma frequently occurs after mastectomy, lymph node dissection, reconstructive procedures, abdominal wall surgery, and abdominoplasty^3–7^.

Seroma formation is generally attributed to disruption of lymphatic and vascular channels, inflammatory exudation, mechanical shear between tissue planes, and persistence of postoperative dead space^8–10^. However, seroma may represent more than a passive fluid collection. Evidence suggests that its development is accompanied by an active tissue response involving inflammatory-cell recruitment, macrophage activity, fibroblast and myofibroblast-associated remodeling, extracellular-matrix deposition, and changes in the local vasculature^11–13^. The temporal relationship between seroma fluid resolution and recovery of the surrounding stromal tissue nevertheless remains poorly defined.

Animal models are essential for investigating postoperative complications and evaluating preventive strategies. However, existing models are heterogeneous and predominantly involve breast, axillary, or donor-site procedures. In a systematic review of 55 surgically induced seroma studies, rats accounted for approximately 69% of the models, and mastectomy for approximately 50% of the surgical procedures^14^. Classical rat seroma models commonly use mastectomy with lymphatic disruption or latissimus dorsi harvest^9,10,15^. In contrast, dedicated abdominal subcutaneous seroma models in rats have rarely been reported, despite the high clinical relevance of abdominal seroma, particularly after abdominoplasty^3,4,7,14^.

Moreover, most previous animal studies have focused on reducing seroma incidence or volume rather than characterizing the biological changes accompanying spontaneous seroma formation and resolution^9,10,14^. In particular, oxidative injury within seroma-associated tissue remains poorly characterized. This represents an important knowledge gap because sustained inflammatory activity can disrupt local redox homeostasis, while oxidative injury may contribute to fibroblast activation, extracellular-matrix remodeling, and persistent inflammation. Lipid peroxidation products such as 4-hydroxynonenal (4-HNE) and oxidized DNA products such as 8-hydroxy-2′-deoxyguanosine (8-OHdG) are established indicators of oxidative tissue injury^16,17^. These processes may interact with macrophage recruitment, collagen deposition, stromal-cell proliferation, and vascular remodeling to prolong tissue pathology after the fluid collection has resolved^18–22^.

Here we describe the development and characterization of a rat post-abdominoplasty seroma model created by controlled separation of the abdominal subcutaneous tissue from the underlying muscle fascia. We monitored seroma formation and volume longitudinally using ultrasound and evaluated seroma-associated tissues at postoperative days (PODs) 14 and 35. Analyses focused on collagen deposition and fiber organization, macrophage-associated inflammatory and reparative responses, lipid and DNA oxidative injury, cellular proliferation, α-smooth muscle actin (α-SMA) expressing stromal and vascular-associated cells, and vascular density. We hypothesized that seroma-associated stromal pathology would persist beyond macroscopic resolution of the fluid collection, with sustained inflammatory, oxidative, fibrotic, and vascular remodeling during the later postoperative phase.

## Results

### Development of the rat abdominal seroma model and longitudinal evolution of seroma volume

To establish a rat model of post-abdominoplasty seroma, the abdominal surgical area was shaved, prepared, and measured to define the operative region (**Supplementary Fig. S1a, b**). We created a transverse incision in the lower abdomen and marked and measured the boundaries of the planned subcutaneous flap (**Supplementary Fig. S1c–f**). The subcutaneous tissue was then elevated from the underlying abdominal muscle fascia, creating a defined dead space between the abdominal soft tissue and muscle layer, followed by closure of the skin incision (**Supplementary Fig. S1g, h**; **Fig. 1a, b**). After surgery, localized swelling progressively developed within the dissected region in animals that subsequently developed seromas (**Fig. 1c**).

**Figure 1.**
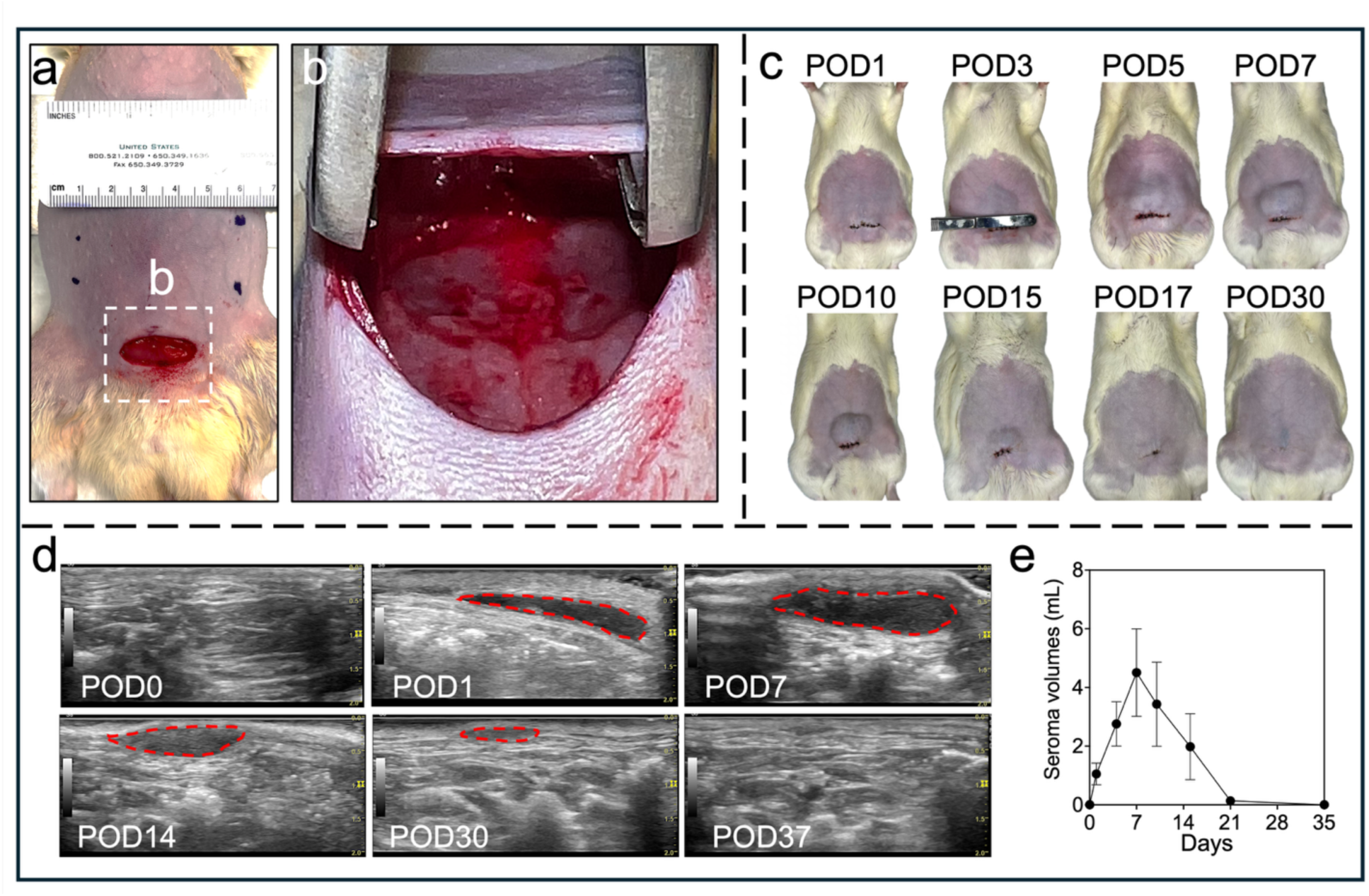
Establishment and longitudinal characterization of the rat post-abdominoplasty seroma model. (a) Preparation of the abdominal surgical field and creation of the transverse incision. The dashed box identifies the region shown in panel b. (b) Representative intraoperative image of the elevated abdominal flap and underlying wound bed following creation of the dead space. (c) Representative gross images of the surgical site from POD1 through POD30. (d) Serial ultrasound images obtained from POD0 through POD37. Red dashed outlines indicate the sonographically identified seroma cavity. (e) Quantification of seroma volume over time, showing fluid accumulation with a peak near POD 7 and subsequent resolution. Data are presented as mean ± SEM. (n = 9)

A total of 20 rats underwent the seroma-induction procedure. Ultrasound-confirmed seroma developed in 9 rats, corresponding to an overall formation rate of 45.0%. The remaining 11 operated rats showed no detectable fluid collection throughout the observation period and were classified as seroma-negative operated animals (**Supplementary Fig. S2**).

Serial ultrasound imaging demonstrated the formation and spontaneous resolution of the seroma cavity (**Fig. 1d**). No fluid collection was detected immediately after surgery (POD0). By POD1, a hypoechoic fluid-filled cavity was visible beneath the abdominal subcutaneous tissue. The cavity progressively enlarged during the first postoperative week, reaching its greatest size at approximately POD7. It remained clearly detectable at POD14 but decreased substantially thereafter. Only minimal residual fluid was observed by POD30, and the cavity was no longer detectable at the final assessment. Quantitative ultrasound analysis confirmed a temporal course characterized by early fluid accumulation followed by progressive spontaneous resolution (**Fig. 1e**). Mean seroma volume increased from approximately 1.1 mL at POD1 to 2.8 mL at POD3 and reached a maximum of approximately 4.5 mL at POD7. It subsequently decreased to approximately 3.4 mL at POD10 and 2.0 mL at POD14. Only minimal residual fluid remained by POD21, and seroma volume approached zero by POD37. Collectively, these findings demonstrate that the procedure produced ultrasound-confirmed abdominal seroma in 45.0% of operated rats, with fluid accumulation peaking during the first postoperative week and resolving spontaneously over the subsequent weeks.

### Seroma formation induces marked and progressive collagen deposition in the abdominal tissue

Masson’s trichrome staining was performed to evaluate stromal collagen deposition in normal abdominal tissue and seroma-associated tissues collected at POD14 and POD35 (**Fig. 2a–i**). Normal tissue exhibited a thin, organized collagen layer between the subcutaneous tissue and underlying muscle (**Fig. 2a–c**). In contrast, POD14 seroma-associated tissue showed marked collagen accumulation and thickening within the surgically created tissue interface (**Fig. 2d–f**). Extensive collagen-rich tissue remained evident at POD35 and appeared denser and thicker than that observed at POD14 (**Fig. 2g–i**).

**Figure 2.**
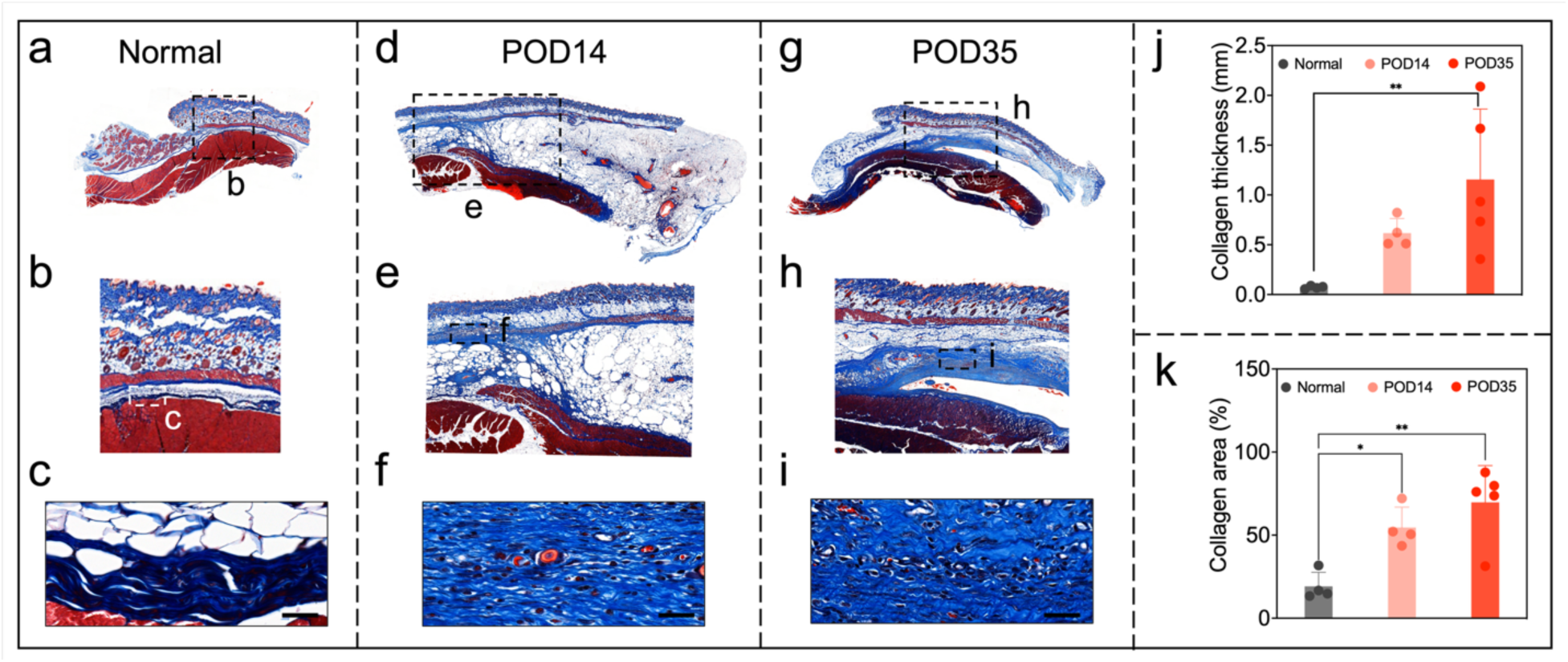
Collagen deposition and capsule thickening after seroma formation. Representative Masson’s trichrome-stained sections from normal tissue (a-c), POD14 tissue (d-f), and POD35 tissue (g-i). Panels a, d, and g show low-magnification views of the tissue architecture. Panels b, e, and h show enlarged views of the boxed regions, and panels c, f, and i show the collagen-rich capsule at higher magnification. Collagen appears blue, muscle and cytoplasm appear red, and nuclei appear dark. (j) Quantification of collagen-layer thickness. (k) Quantification of collagen-positive area. Scale bars: 40 μm. Data are presented as mean ± SEM. Normal tissue (n = 4), POD 14 (n = 4), and POD 35 (n = 5). *P <0.05, **P <0.01, ***P <0.001.

Quantitative analysis showed that the collagen-layer thickness increased from approximately 0.1 mm in normal tissue to 0.6 mm at POD14 and 1.2 mm at POD35, with a significant difference between the normal and POD35 groups (**Fig. 2j**; P <0.01). The collagen-positive area increased from approximately 19% in normal tissue to 54% at POD14 and 69% at POD35 (**Fig. 2k**). Compared with normal tissue, the collagen-positive area was significantly greater at both POD14 (P <0.05) and POD35 (P <0.001). Although both collagen thickness and collagen-positive area were numerically higher at POD35 than at POD14, no significant direct comparison between these time points was demonstrated. These findings indicate that pronounced fibrotic remodeling persists within seroma-associated tissue even as the fluid collection progressively resolves.

Picrosirius red staining was used to evaluate the organization and structural characteristics of collagen fibers in normal abdominal tissue and seroma-associated tissues collected at POD14 and POD35 (Fig. 3a–f). Under polarized light, normal tissue was dominated by red birefringent fibers, consistent with relatively thick, densely packed collagen bundles (Fig. 3a, b). Seroma-associated tissues showed a progressive increase in yellow and green birefringent fibers at POD14 and POD35 (Fig. 3c–f), indicating alterations in collagen fiber thickness and organization.

**Figure 3.**
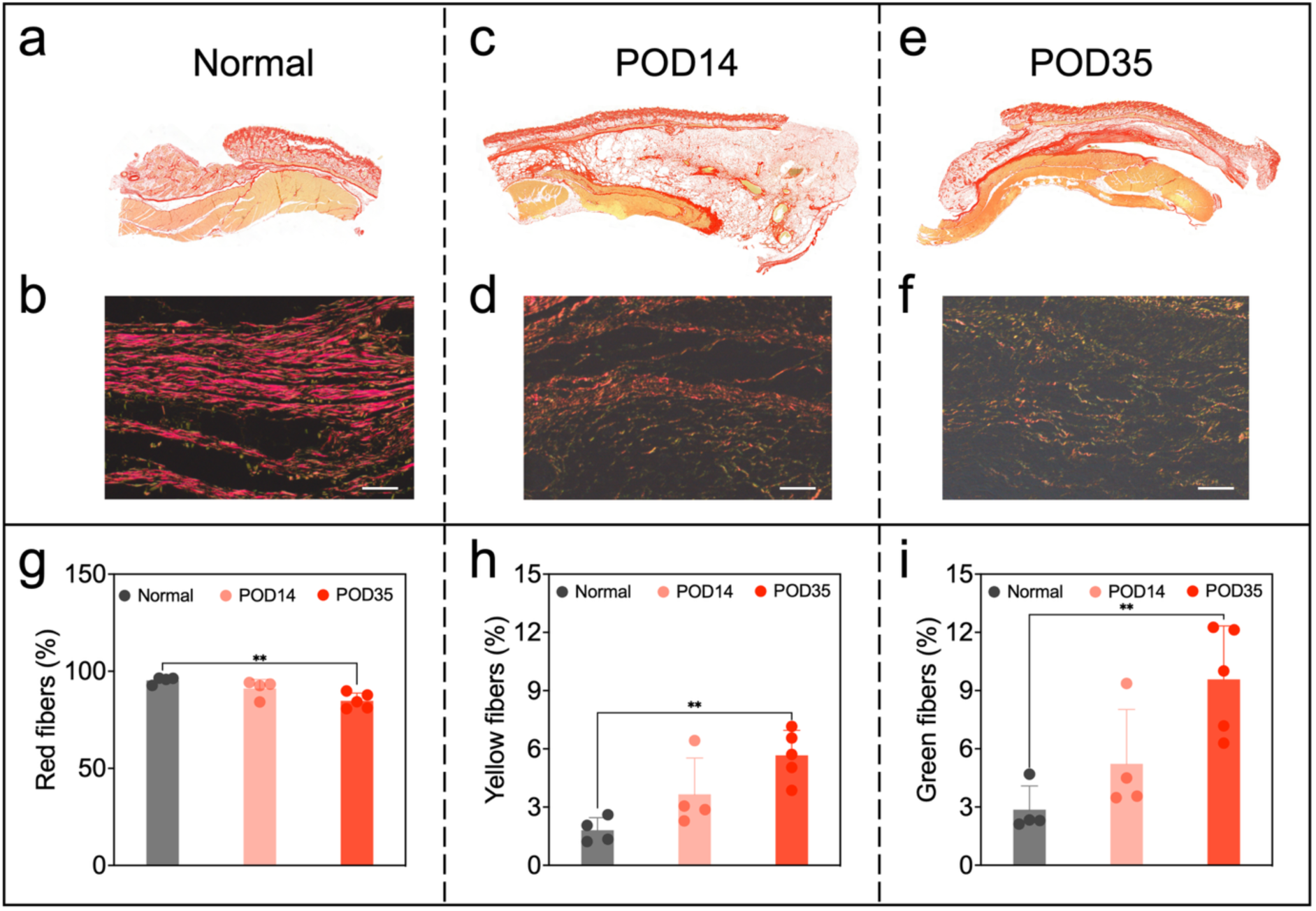
Collagen fiber remodeling during seroma-capsule development. Representative picrosirius red-stained (PSR) sections from normal tissue (a, b), POD14 tissue (c, d), and POD35 tissue (e, f). Panels a, c, and e show bright-field images, and panels b, d, and f show collagen birefringence under polarized light. (g-i) Quantification of red (g), yellow (h), and green (i) birefringent collagen fibers. Scale bars: 50 μm. Red, yellow, and green birefringence should be interpreted as differences in fiber thickness, packing, and organization rather than as direct measures of collagen type. PSR images at POD14 and POD35 were selected from the seroma capsule region. Data are presented as mean ± SEM. Normal tissue (n = 4), POD 14 (n = 4), and POD 35 (n = 5). \**P <*0.05, **P <0.01.

Quantitative analysis showed that the proportion of red fibers decreased from approximately 95% in normal tissue to 91% at POD14 and 85% at POD35, with a significant difference between the normal and POD35 groups (**Fig. 3g**; P <0.01). Conversely, yellow fibers increased from approximately 2% in normal tissue to 4% at POD14 and 6% at POD35 (**Fig. 3h**), while green fibers increased from approximately 3% to 5% and 9%, respectively (**Fig. 3i**). The proportions of both yellow and green fibers were significantly greater at POD35 than in normal tissue (P <0.01). Although the changes were numerically more pronounced at POD35 than at POD14, we did not demonstrate a significant direct comparison between the two postoperative time points. Together with the increased collagen deposition shown by Masson’s trichrome staining, these findings indicate that seroma-associated tissue undergoes sustained collagen accumulation accompanied by persistent remodeling of collagen fiber architecture.

### Persistent macrophage accumulation and mixed inflammatory remodeling in seroma-associated tissue

Immunofluorescence staining was performed to characterize macrophage accumulation and phenotype in normal and seroma-associated tissues (**Fig. 4**). CD68 was used as a pan-macrophage marker, whereas CD86 was used to identify macrophages with a pro-inflammatory phenotype and CD206 was used as a marker associated with a reparative, alternatively activated macrophage phenotype^23,24^. CD68-positive cells were sparse in normal abdominal tissue but markedly increased at POD14 and POD35 (**Fig. 4a**). Both the percentage and density of CD68-positive cells were significantly greater at POD14 and POD35 than in normal tissue (**Fig. 4b, c**). Moreover, CD68-positive cell density increased significantly from POD14 to POD35, indicating continued macrophage accumulation during the later phase of seroma-associated tissue remodeling (**Fig. 4c**).

**Figure 4.**
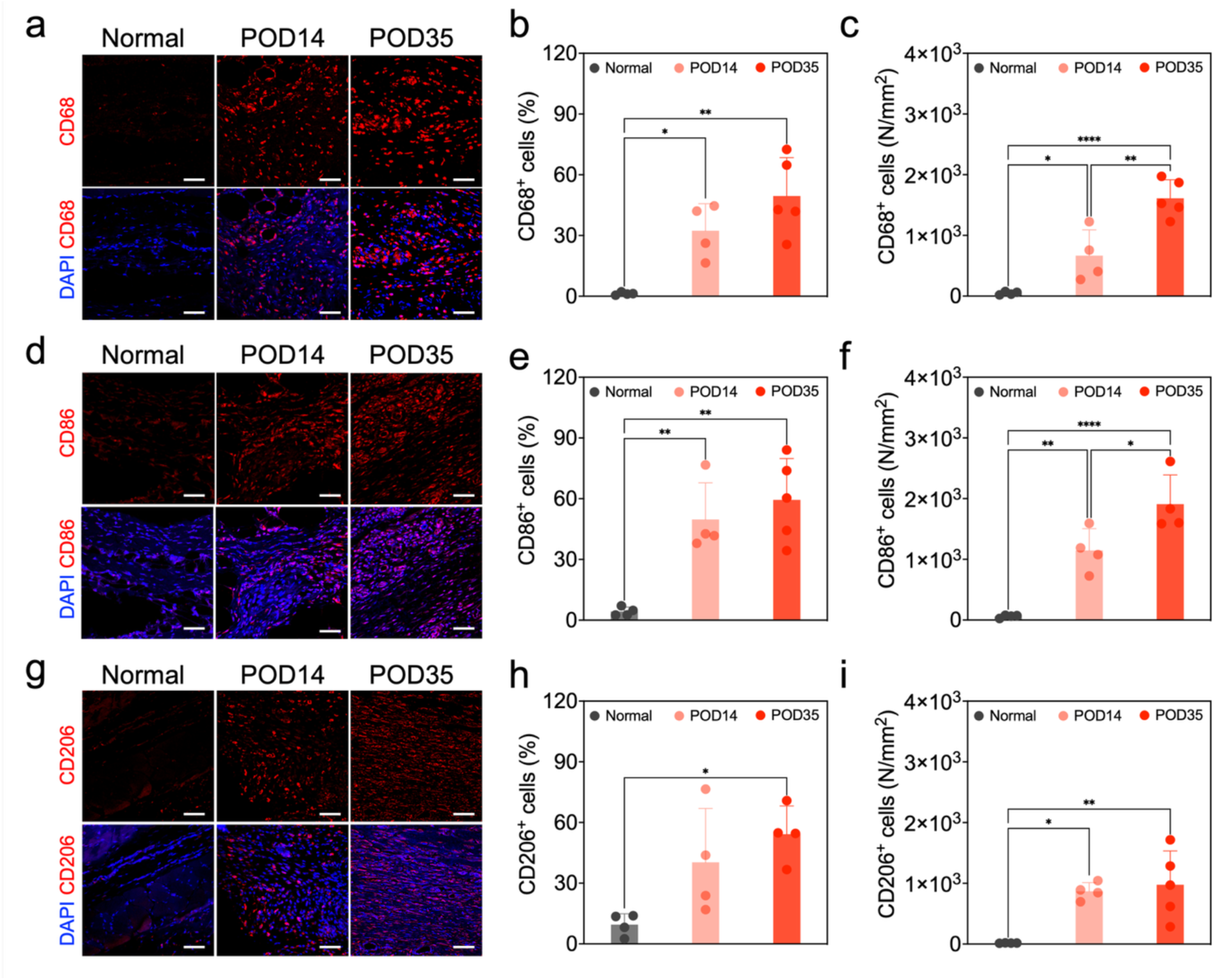
CD68, CD86, and CD206 immunoreactivity during seroma-capsule development. Representative immunofluorescence images and quantitative analyses of CD68 (a-c), CD86 (d-f), and CD206 (g-i) in normal, POD14, and POD35 tissues. Panels a, d, and g show marker staining in red and *nuclear* counterstaining with DAPI in blue. Panels b, e, and h report the percentage of marker-positive cells, and panels c, f, and i report marker-positive cell density per square millimeter. Scale bars: 50 μm. Representative macrophage images at POD14 and POD35 were acquired from the seroma capsule region. Data are presented as mean ± SEM. Normal tissue (n = 4), POD 14 (n = 4), and POD 35 (n = 5). \**P* <0.05, \*\**P* <0.01, \*\*\**P* <0.001, \*\*\*\**P* <0.0001.

CD86-positive cells were also markedly increased in seroma-associated tissue at both postoperative time points (**Fig. 4d**). The percentage and density of CD86-positive cells were significantly greater at POD14 and POD35 than in normal tissue (**Fig. 4e, f**). CD86-positive cell density increased further from POD14 to POD35 (**Fig. 4f**), demonstrating persistence and intensification of a pro-inflammatory cellular response despite progressive resolution of the seroma fluid.

CD206-positive cells were more abundant in seroma-associated tissue than in normal tissue (**Fig. 4g**). The percentage of CD206-positive cells was significantly increased at POD35 relative to normal tissue, while CD206-positive cell density was significantly elevated at both POD14 and POD35 (**Fig. 4h, i**). No significant difference in CD206-positive cell density was detected between POD14 and POD35.

Collectively, these findings demonstrate sustained macrophage accumulation within seroma-associated tissue, accompanied by concurrent increases in CD86- and CD206-associated cellular responses. The further elevation of total CD68-positive and CD86-positive cell densities at POD35 suggests that inflammatory remodeling persists and may intensify even after the seroma fluid has largely resolved.

### Persistent lipid peroxidation and oxidative DNA damage are present during seroma remodeling

To determine whether oxidative stress contributes to the stromal pathology associated with seroma formation, tissue sections were evaluated for 4-hydroxynonenal (4-HNE), a marker of lipid peroxidation, and 8-hydroxy-2′-deoxyguanosine (8-OHdG), a marker of oxidative DNA damage (**Fig. 5**).

**Figure 5.**
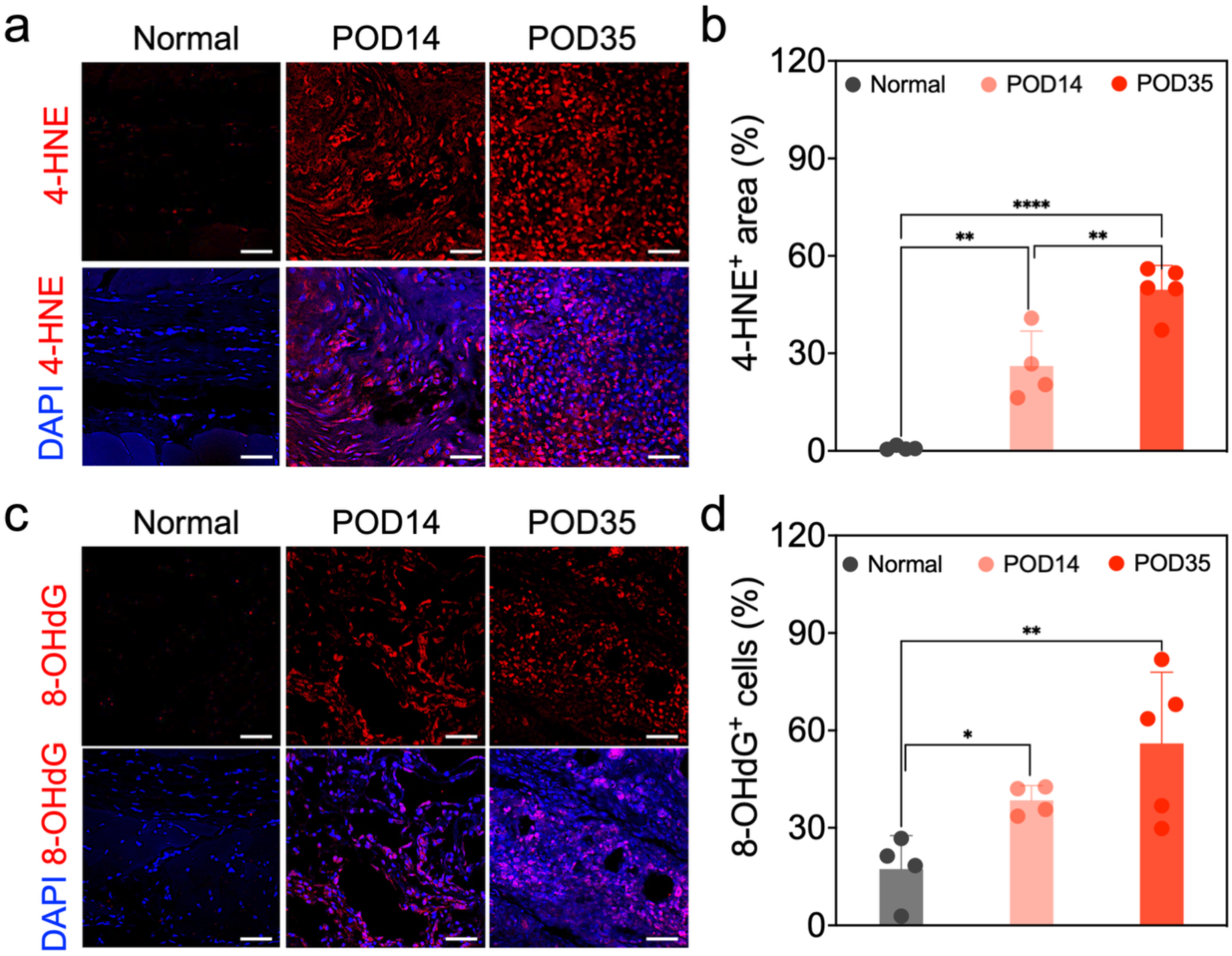
Oxidative damage increases during seroma-capsule development. (a) Representative immunofluorescence images of 4-hydroxynonenal (4-HNE; red), a marker of lipid peroxidation, in normal, POD14, and POD35 tissues. Nuclei were counterstained with DAPI (blue). (b) Quantification of 4-HNE-positive area. (c) Representative immunofluorescence images of 8-hydroxy-2’-deoxyguanosine (8-OHdG; red), a marker of oxidative DNA damage, in normal, POD14, and POD35 tissues. (d) Quantification of 8-OHdG-positive cells. Scale bars: 50 μm. Representative oxidative damage images at POD14 and POD35 were acquired from the seroma capsule region. Data are presented as mean ± SEM. Normal tissue (n = 4), POD 14 (n = 4), and POD 35 (n = 5). \**P* <0.05, \*\**P* <0.01, \*\*\*\**P* <0.0001.

Minimal 4-HNE immunoreactivity was detected in normal abdominal tissue, whereas markedly increased staining was observed in seroma-associated tissues at POD14 and POD35 (**Fig. 5a**). The 4-HNE-positive area increased from approximately 1% in normal tissue to 26% at POD14 and 49% at POD35 (**Fig. 5b**). Compared with normal tissue, 4-HNE staining was significantly elevated at both POD14 (P <0.01) and POD35 (P <0.0001). Importantly, the 4-HNE-positive area was also significantly greater at POD35 than at POD14 (P <0.01), demonstrating progressive lipid peroxidation during the later phase of tissue remodeling.

Similarly, 8-OHdG-positive cells were increased in seroma-associated tissue at both postoperative time points (**Fig. 5c**). The percentage of 8-OHdG-positive cells rose from approximately 17% in normal tissue to 38% at POD14 and 56% at POD35 (Fig. 5d). These values were significantly higher than normal at POD14 (P <0.05) and POD35 (P <0.01). Although the percentage was numerically greater at POD35 than at POD14, no significant difference was observed between the two postoperative time points.

Together, these findings reveal persistent oxidative injury within seroma-associated tissue. In particular, lipid peroxidation continued to increase from POD14 to POD35, while oxidative DNA damage remained elevated, despite the near-complete resolution of the seroma fluid. This temporal dissociation suggests that macroscopic fluid resolution does not indicate recovery of the underlying stromal microenvironment.

### Proliferative stromal and vascular responses peaked during the early remodeling phase

Ki67 and α-smooth muscle actin (αSMA) co-staining was performed to assess cellular proliferation and the involvement of αSMA-expressing stromal and vascular cells during seroma-associated tissue remodeling (Fig. 6a). Normal abdominal tissue contained few Ki67-positive cells, whereas POD14 showed a marked increase. Ki67-positive cells remained detectable at POD35 but were less abundant than at POD14. High-magnification images demonstrated Ki67 and αSMA colocalization in stromal cells with myofibroblast-like morphology and in cells associated with vascular structures (**Fig. 6b**).

**Figure 6.**
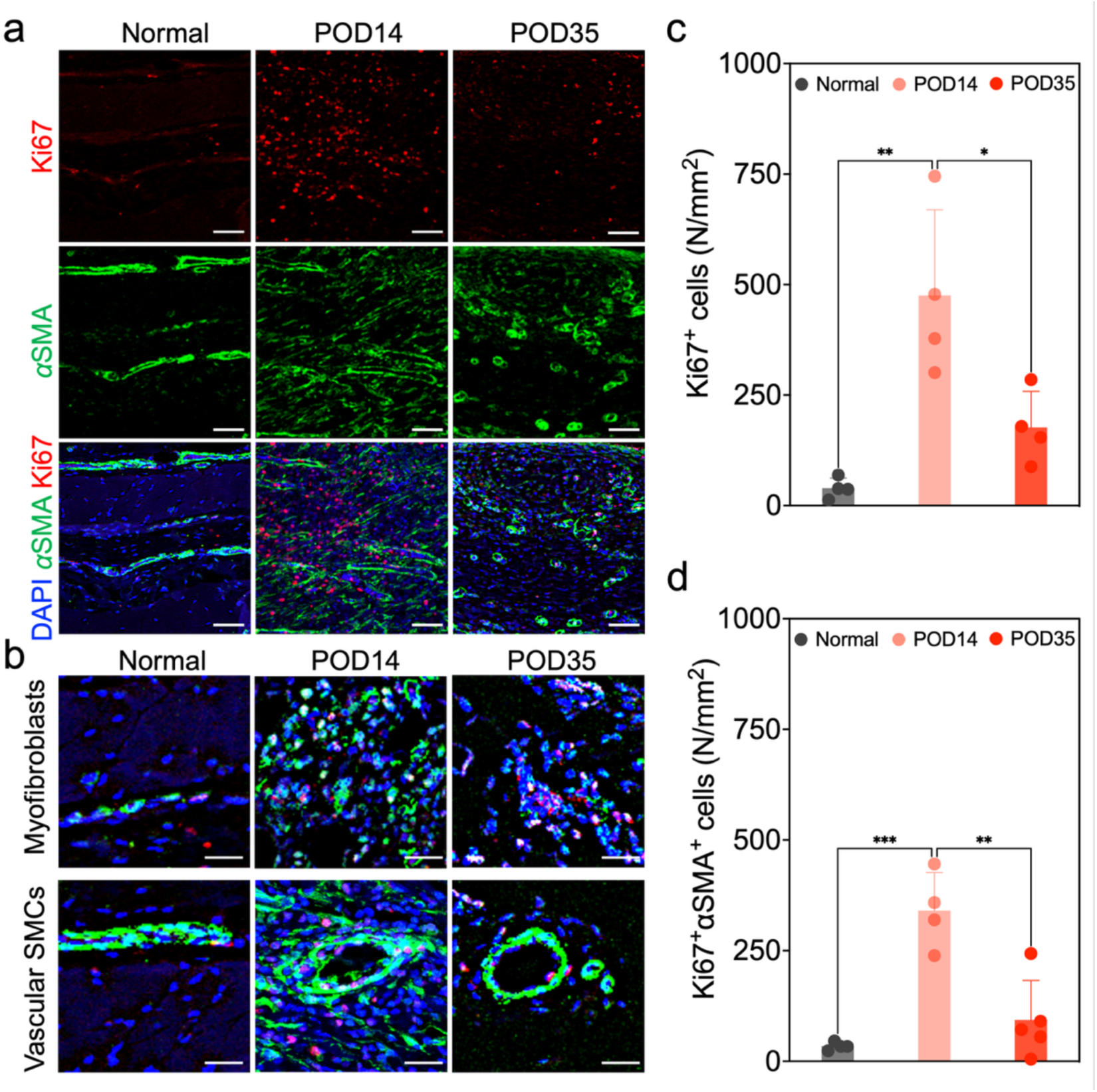
Cell proliferation and proliferative and αSMA-associated cellular responses during capsule remodeling. (a) Representative immunofluorescence images of Ki67 (red) and αSMA (green) in normal, POD14, and POD35 tissues. Nuclei were counterstained with DAPI (blue). (b) Higher-magnification images showing proliferating αSMA-positive myofibroblasts and αSMA-positive vascular smooth muscle cells. (c) Quantification of Ki67-positive cell density. (d) Quantification of Ki67 and αSMA double-positive cell density. Scale bars: 50 μm. Representative immunofluorescence images at POD14 and POD35 were obtained from the seroma capsule region. Data are presented as mean ± SEM. Normal tissue (n = 4), POD 14 (n = 4), and POD 35 (n = 5). \**P* <0.05, \*\**P* <0.01, \*\*\**P* <0.001.

Quantitative analysis showed that Ki67-positive cell density increased from approximately 40 cells/mm² in normal tissue to 475 cells/mm² at POD14 (P <0.01; **Fig. 6c**). By POD35, Ki67-positive cell density had decreased to approximately 176 cells/mm² and was significantly lower than at POD14 (P <0.05). Similarly, the density of Ki67 and αSMA double-positive cells increased from approximately 34 cells/mm² in normal tissue to 340 cells/mm² at POD14 (P <0.001; **Fig. 6d**), before decreasing to approximately 93 cells/mm² at POD35 (P <0.01 versus POD14).

These findings demonstrate a robust but temporally restricted proliferative response involving αSMA-expressing stromal and vascular-associated cells. This response peaked at POD14 and subsequently declined, indicating a transition from an early proliferative phase toward a later phase characterized by persistent fibrosis, inflammation, and oxidative injury.

### Increased vascular density persisted during late seroma-associated tissue remodeling

To evaluate vascular remodeling within seroma-associated tissue, sections were co-stained for the endothelial marker CD31 and α-smooth muscle actin (αSMA) (Fig. 7). In contrast, vascular structures were markedly more abundant within seroma-associated tissue at both POD14 and POD35 (**Fig. 7a**). Many CD31-positive vessels were surrounded by αSMA-positive cells, demonstrating the presence of endothelialized vessels with associated mural-cell coverage. Quantitative analysis showed that vessel density increased from approximately 12 vessels/mm² in normal tissue to 49 vessels/mm² at POD14 and 64 vessels/mm² at POD35 (**Fig. 7b**). Compared with normal tissue, vessel density was significantly greater at POD14 (P <0.05) and POD35 (P <0.01). Although vessel density was numerically higher at POD35 than at POD14, no significant difference was demonstrated between the two postoperative time points. These findings demonstrate sustained vascular remodeling within seroma-associated tissue. Notably, increased vascular density persisted at POD35 despite the decline in cellular proliferation and near-complete resolution of the seroma fluid, indicating that the remodeled stromal microenvironment had not returned to its normal vascular state.

**Figure 7.**
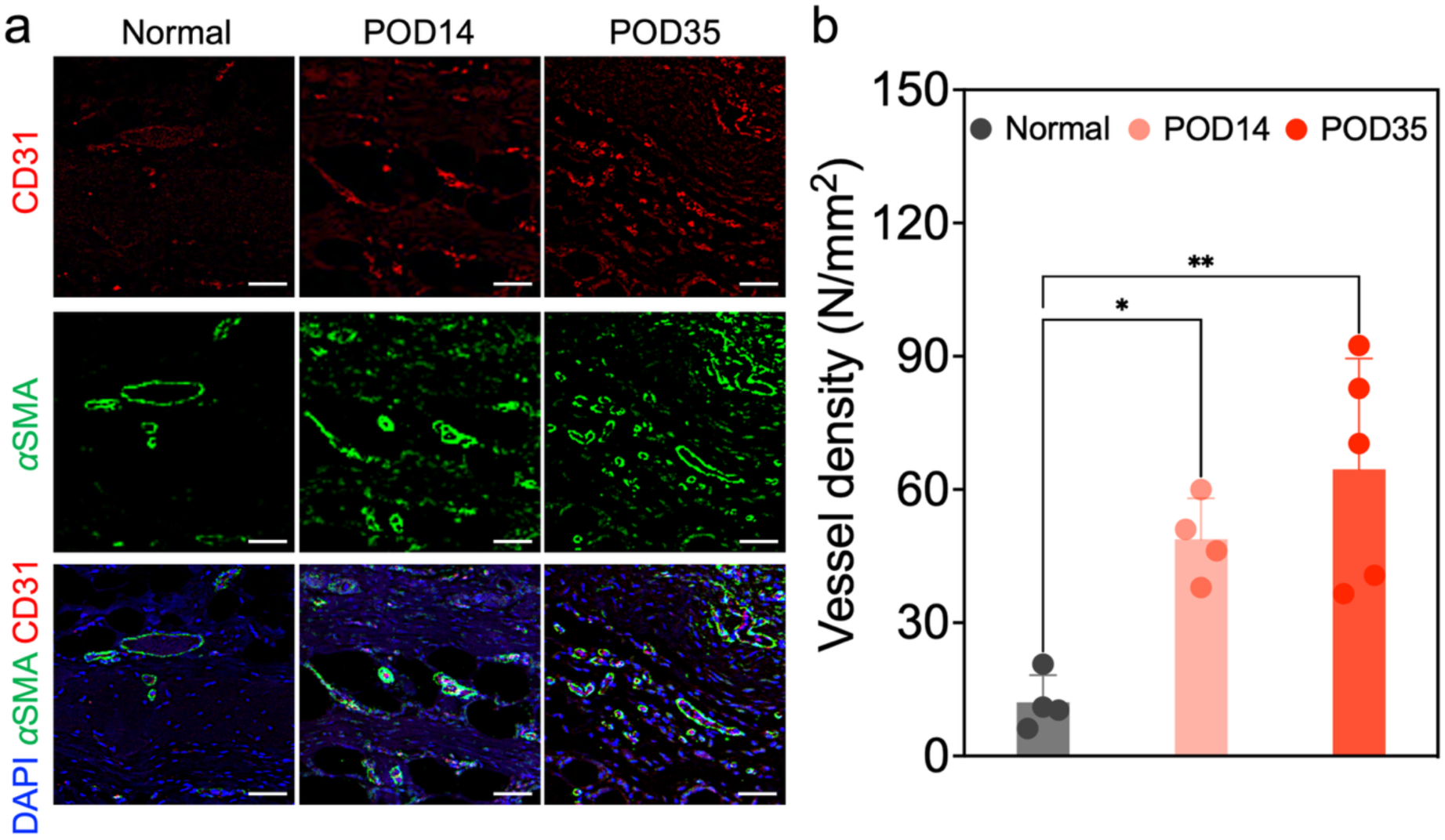
Vascular density increases during seroma-capsule remodeling. (a) Representative immunofluorescence images of CD31 (red) and αSMA (green) in normal, POD14, and POD35 tissues. Nuclei were counterstained with DAPI (blue). (b) Quantification of vessel density per square millimeter. Scale bars: 50 μm. Representative images of blood vessels at POD14 and POD35 were obtained from the seroma capsule region. Data are presented as mean ± SEM. Normal tissue (n = 4), POD 14 (n = 4), and POD 35 (n = 5). \**P* <0.05, \*\**P* <0.01.

## Discussion

The principal finding of this study is that macroscopic resolution of postoperative seroma does not coincide with biological recovery of the surrounding stromal tissue. We developed and longitudinally characterized a rat abdominal subcutaneous seroma model that reproduced both postoperative fluid accumulation and a prolonged tissue-remodeling response. Although experimental seroma models have been investigated for several decades, previous studies have predominantly used mastectomy, axillary lymphatic disruption, or latissimus dorsi harvest models^9,10,14,15^. In contrast, we created the present model by elevating the abdominal subcutaneous tissue from the underlying muscle fascia, reproducing the broad tissue-plane separation and dead space generated during abdominoplasty. To our knowledge, this is the first rat model specifically designed to reproduce post-abdominoplasty seroma and to characterize the temporal evolution of oxidative stress within seroma-associated tissue.

Ultrasound-confirmed seroma developed in 9 of 20 operated animals, corresponding to a formation rate of 45.0%. Fluid accumulation became detectable during the early postoperative period, reached a maximum volume of approximately 5 mL at POD7, and then spontaneously resolved, approaching zero by POD21. Thus, controlled separation of the abdominal subcutaneous tissue from the muscle fascia was sufficient to induce a measurable and self-resolving seroma in approximately half of the operated animals without mastectomy, lymphadenectomy, or muscle resection. The incomplete formation rate may reflect biological variability in the response to soft-tissue dissection and dead-space creation. This characteristic also resembles the clinical setting, in which seroma develops in only a subset of patients undergoing similar surgical procedures^3^.

Importantly, the model reproduced not only transient fluid accumulation but also a prolonged tissue-remodeling response. Despite near-complete resolution of the seroma fluid by POD35, substantial biological alterations remained evident. Collagen deposition progressively increased, with collagen thickness rising from approximately 0.1 mm in normal tissue to 0.6 mm at POD14 and 1.2 mm at POD35, while collagen-positive area increased from approximately 19% to 54% and 69%, respectively. Macrophage accumulation was similarly pronounced, with CD68+ cells increasing from approximately 1.3% in normal tissue to 32.3% at POD14 and 49.5% at POD35. Vessel density also increased progressively from approximately 12 vessels/mm² in normal tissue to 49 vessels/mm² at POD14 and 64 vessels/mm² at POD35. Notably, oxidative injury persisted throughout the remodeling period. Both 4-HNE, a marker of lipid peroxidation, and 8-OHdG, a marker of oxidative DNA damage, remained markedly elevated at POD35^16,17^, when the seroma cavity was nearly undetectable. This temporal dissociation between fluid resolution and persistent oxidative injury indicates that seroma disappearance does not represent complete restoration of the local tissue environment. Instead, sustained oxidative stress accompanies ongoing fibrotic, inflammatory, and vascular remodeling during the later stages of seroma resolution. However, a limitation to our analyses is an even further endpoint, i.e. >POD 35, to make further determination as to whether fibrotic and inflammatory remodeling fully resolves given sufficient time or is in a chronic remodeling state.

This model may therefore be particularly useful for investigating seroma following abdominal soft-tissue procedures, in which broad tissue-plane separation and persistent dead space contribute to postoperative fluid accumulation^2,14,25^. The model permits longitudinal ultrasound monitoring of seroma kinetics together with terminal characterization of the surrounding stromal response. Clinically, the findings suggest that disappearance or drainage of seroma fluid may not indicate complete normalization of the affected tissue. Strategies that target fluid accumulation alone may therefore differ from approaches intended to modulate the persistent inflammatory, oxidative, and fibrotic microenvironment. The model provides a platform to test dead-space closure methods, tissue adhesives, anti-inflammatory or antioxidant interventions, and other strategies aimed at both seroma prevention and pathological stromal remodeling.

These findings should be interpreted in the context of several considerations that also guide future investigation. The present histological analyses compared seroma-positive tissues with nonsurgical normal tissues and identified substantial tissue changes associated with the postoperative seroma environment. Future inclusion of seroma-negative operated controls will further distinguish the specific effects of seroma formation from those of surgical dissection and dead-space creation. Seroma developed in 45.0% of operated animals, reflecting clinically relevant variability while highlighting the opportunity to further optimize model consistency for intervention studies. The 4-HNE and 8-OHdG findings establish oxidative injury as a prominent feature of seroma-associated tissue remodeling, although additional studies are needed to identify the cellular sources of reactive oxygen species and determine their causal contributions to fibrosis and inflammation. Finally, the two terminal time points define distinct early and established stages of seroma-associated remodeling. Building on this foundation, additional temporal assessments and targeted antioxidant or anti-inflammatory interventions will help clarify the mechanisms linking oxidative stress to persistent stromal remodeling.

## Conclusion

We developed and characterized a rat model that reproduces seroma formation within the post-abdominoplasty abdominal tissue plane, with ultrasound-confirmed seroma developing in 45.0% of operated animals. Seroma volume peaked during the first postoperative week and resolved spontaneously over approximately 5 weeks. However, fluid collection resolution did not coincide with normalization of the surrounding stromal tissue. Seroma-associated tissue exhibited persistent collagen accumulation and remodeling, macrophage infiltration, oxidative lipid and DNA injury, and increased vascular density. The cellular proliferative response peaked at POD14 and subsequently declined, whereas fibrosis, inflammatory cell accumulation, oxidative injury, and vascular remodeling remained evident at POD35. Notably, lipid peroxidation and the densities of CD68-positive and CD86-positive cells increased further between POD14 and POD35. These findings suggest a temporal dissociation between macroscopic seroma resolution and biological stromal recovery through POD35. This model provides a platform for investigating post-abdominoplasty seroma pathogenesis and suggests that therapeutic strategies may need to consider not only fluid accumulation but also the inflammatory, oxidative, and fibrotic tissue alterations that persist after macroscopic seroma resolution.

## Materials and Methods

### Animal Use and Ethical Statement

All animal procedures were approved by the Institutional Animal Care and Use Committee of Washington University in St. Louis (Protocol no. 23-0313) and were conducted in accordance with institutional guidelines. The study is reported in accordance with the ARRIVE 2.0 guidelines. Male Sprague–Dawley rats [Crl:CD(SD), strain code 001] were purchased from Charles River Laboratories (Wilmington, MA, USA), weighed approximately 400 g at the time of surgery, and were housed under standard laboratory conditions with ad libitum access to food and water. For surgery, rats were anesthetized with inhaled isoflurane (1–3% in oxygen via a precision vaporizer) and subsequently administered ketamine/dexmedetomidine (75 mg/kg and 0.5 mg/kg, intraperitoneally). Adequate anesthetic depth was confirmed by the absence of the pedal withdrawal reflex and by physiological monitoring throughout the procedure. Analgesia was provided with sustained-release buprenorphine (0.5 mg/kg, subcutaneously; a single administration providing up to 72 h of analgesia). At the designated study endpoints, animals were euthanized by intracardiac injection of Fatal-Plus® (pentobarbital sodium, 390 mg/mL; 100 µL per rat). Death was confirmed by cessation of cardiac activity and respiration. Of the 20 rats that underwent the seroma-induction procedure, ultrasound-confirmed seroma developed in nine animals. Among the seroma-positive animals, tissues were collected at POD14 (n = 4) or POD35 (n = 5) for histological and immunofluorescence analyses, representing active and late post-resolution phases of seroma-associated tissue remodeling, respectively. Four additional rats served as nonsurgical normal controls, from which normal abdominal tissues were collected (n = 4). The remaining 11 operated rats did not develop ultrasound-detectable seroma and were classified as seroma-negative operated animals. These animals were included in the assessment of seroma incidence but were not included in the terminal tissue analyses.

### Rat Abdominal Seroma Model

Rats were anesthetized with inhaled isoflurane, and the abdominal hair was removed. Following aseptic preparation, a 3-cm transverse incision was created in the lower abdominal region. A 6 × 6 cm dissection field extending cranially from the incision was measured and marked. Through the incision, the skin and subcutaneous tissue were elevated from the underlying abdominal muscle fascia within the marked boundaries, creating a defined subcutaneous dead space while preserving the muscle fascia. The incision was subsequently closed with interrupted sutures. No drain was placed, and the dead space was not otherwise obliterated. Animals received postoperative analgesia according to the approved institutional protocol and were monitored for wound healing, surgical complications, and seroma formation. The sequential operative procedure is illustrated in **Supplementary Fig. S1**.

### Ultrasound Assessment of Seroma

Seroma formation and resolution were monitored longitudinally by ultrasound imaging. The seroma cavity was identified as a hypoechoic fluid-containing space between the abdominal subcutaneous tissue and the underlying muscle fascia. Ultrasound imaging was performed at selected postoperative time points from POD0 through POD35. Seroma volume was quantified from ultrasound images and used to evaluate the temporal progression and spontaneous resolution of the fluid collection. Seroma-positive animals were defined as those with a measured seroma volume greater than 0.2 mL within the surgically created subcutaneous dead space. Nine rats met this criterion. The remaining 11 operated rats showed no detectable seroma throughout the observation period and were classified as seroma-negative (**Supplementary Fig. S2**). Seroma became detectable during the early postoperative period, reached its maximum volume around POD7, and gradually decreased thereafter, becoming nearly undetectable by POD30–35.

### Histology and Collagen Analysis

At designated endpoints, abdominal tissues containing the seroma region and surrounding remodeled tissue were harvested, fixed in 10% neutral-buffered formalin, paraffin-embedded, and sectioned at 5 μm thickness. Hematoxylin and eosin (H&E) staining was performed to evaluate general tissue morphology and structural remodeling. Masson’s trichrome staining was used to assess collagen deposition and fibrosis. Collagen thickness and collagen-positive area were quantified using ImageJ/Fiji from representative regions of each tissue section. Picrosirius Red (PSR) staining combined with polarized light microscopy was used to evaluate collagen fiber organization^26^. Collagen fibers were classified according to birefringent color as red/orange, yellow, or green, and the relative percentage of each fiber population was quantified using ImageJ/Fiji.

### Immunofluorescence Analysis

Paraffin sections were deparaffinized, rehydrated, and subjected to antigen retrieval. Following blocking, sections were incubated overnight at 4°C with the appropriate primary antibodies. Macrophage accumulation and phenotype were evaluated using CD68, CD86, and CD206 staining. Positive cells were quantified as either percentage of total cells or cell density within the analyzed tissue area. Cellular proliferation and αSMA-associated remodeling were assessed by Ki67 and αSMA co-staining. Ki67-positive cells and Ki67+/αSMA+ cells were quantified within representative regions of the remodeled tissue. Vascular remodeling was evaluated by CD31 and αSMA co-staining. Vessel density was quantified as the number of CD31-positive vascular structures per mm². Oxidative tissue injury was evaluated by immunofluorescence staining for 4-hydroxynonenal (4-HNE) and 8-hydroxy-2′-deoxyguanosine (8-OHdG). 4-HNE was used as a marker of lipid peroxidation, whereas 8-OHdG was used as a marker of oxidative DNA damage^17^. Positive staining was quantified using ImageJ/Fiji and expressed as the percentage of positive area or positive cells within standardized regions of interest.

### Image Analysis

Image analysis included collagen thickness and area for Masson’s trichrome; red-orange, yellow, and green birefringent collagen fibers for polarized picrosirius red (PSR); CD68⁺, CD86⁺, and CD206⁺ cell density per mm²; 4-HNE⁺ area and 8-OHdG⁺ cell density; Ki67⁺ and Ki67⁺/α-SMA⁺ cell density; and vessel density per mm².

All images were analyzed in Fiji/ImageJ. Processing parameters were applied uniformly to all images within experimental datasets. For fluorescence images, the red, green, and blue channels were separated using the Split Channels function and converted to 8-bit grayscale images. Background fluorescence was corrected using Subtract Background with a predefined rolling-ball radius. Intensity thresholds were determined empirically for each channel to distinguish specific signal from background and once established, were kept unchanged across all images in the corresponding dataset. When necessary, regions of interest were drawn to limit the analysis to the relevant tissue compartment. Image dimensions were calibrated using the acquisition scale.

For area-based outcomes, including collagen⁺ area and 4-HNE⁺ area, the thresholded signal area was measured relative to the total tissue area analyzed. For polarized PSR, RGB images were analyzed after conversion to hue, saturation, and brightness (HSB) components. Birefringent collagen fibers were classified using predefined 8-bit hue ranges: red-orange, 0–38 and 230–255; yellow, 39–51; and green, 52–128, with a saturation threshold of ≥40 and a brightness threshold of ≥20. Binary masks were generated for red-orange, yellow, and green fibers, and their sum was used to define total birefringent collagen fiber area. Segmentation masks and overlays were reviewed during analysis to confirm appropriate signal detection and to identify obvious artifacts.

For nuclear and cell-based measurements, the DAPI channel was thresholded to generate a binary nuclear mask. Adjacent nuclei were separated using the Watershed function, after which individual objects were identified with Analyze Particles. Predefined particle-size criteria were used to exclude debris and background. Marker⁺ cells were identified based on segmentation mask overlaps. For measurements with multiple positive markers, including Ki67⁺/α-SMA⁺ cells, the corresponding segmentation masks were spatially compared. For density-based outcomes, including cell density and vessel density, analyzed tissue areas were converted to mm², and density was calculated as the number of positive marker objects divided by the analyzed area in mm².

### Statistical Analysis

Data are presented as means ± standard error of the mean (SEM). Statistical analyses were performed in GraphPad Prism 10. Outcomes included collagen-layer thickness, collagen-positive area, red-orange, yellow, and green birefringent collagen fibers, CD68⁺, CD86⁺, and CD206⁺ cell density, 4-HNE⁺ area, 8-OHdG⁺ cell density, Ki67⁺ cell density, Ki67⁺/α-SMA⁺ double-positive cell density, and vessel density. When multiple fields were obtained from the same biological specimen, measurements were averaged to generate a single value for that specimen before statistical testing. We compared the Normal, POD14, and POD35 groups using an ordinary one-way ANOVA followed by Tukey’s multiple-comparisons test. All tests were two-sided, and adjusted *P* values < 0.05 were considered statistically significant. Statistical significance is indicated as follows: \**P <*0.05, \*\**P <*0.01, \*\*\**P <*0.001, and \*\*\*\**P <*0.0001.

## Data Availability

The datasets generated and analyzed during the current study are available from the corresponding authors upon reasonable request.

## Author Contributions

X.X. contributed to the design, execution, and analysis of all experiments. A.M.N., K.H., A.B., and Y.G. performed experiments and contributed to data analysis. S.H., B.A., and L.S. assisted with sample preparation, histology, and image quantification. A.E.G., M.A.Z., and M.W. provided critical input on experimental design, interpretation of the findings, and manuscript revision. J.M.S. and X.L. conceptualized the project, supervised all aspects of the study, and interpreted the results. All authors reviewed and approved the final manuscript.

## Acknowledgements

This work was supported by the Center of Regenerative Medicine (CRM) Senior Innovation Award from Washington University School of Medicine and National Institutes of Health grant R01HL168513 (X.L.).

