## Supplemental Figures 1 and 2 for "Oxidative Stress and Tissue Remodeling in a Rat Abdominal Seroma Model"

Corresponding Authors:

Justin M. Sacks, MD

### Supplementary Information

#### Figure S1

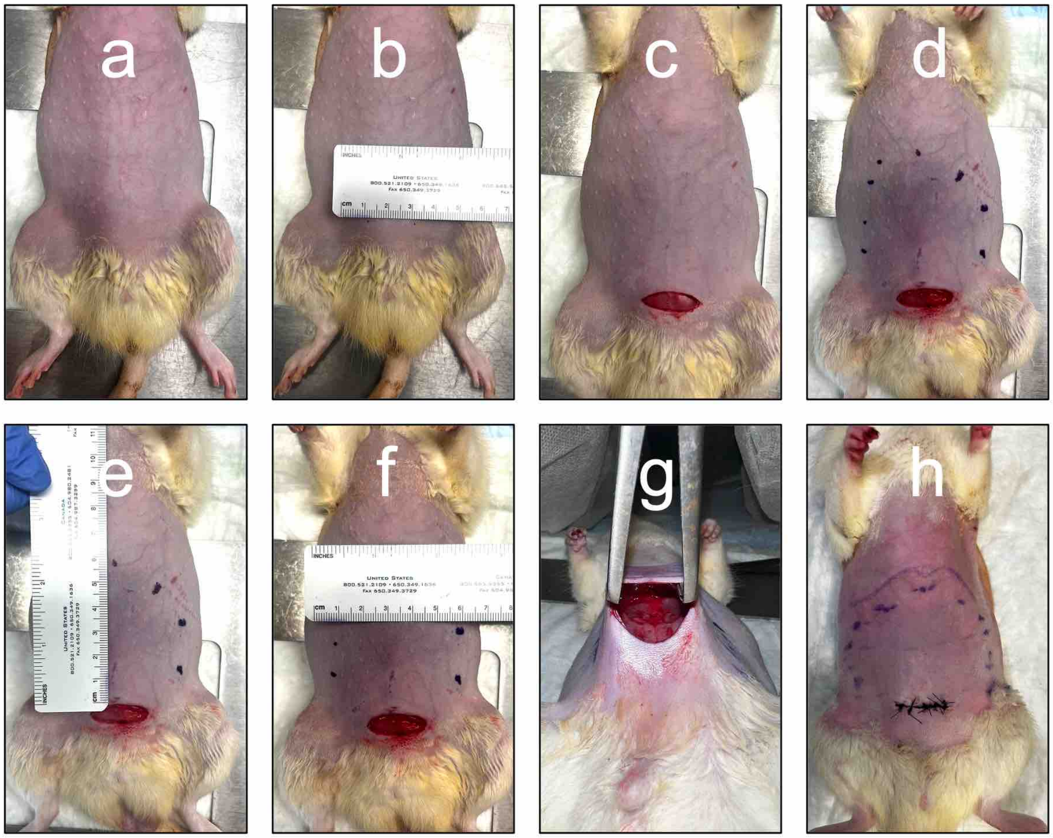


**Figure S1. Surgical procedure used to establish the rat post-abdominoplasty seroma model.** Representative sequential photographs of the operative procedure: preparation of the abdominal field (a), measurement and marking of the operative area (b-f), elevation of the abdominal flap and creation of the wound bed (g), and closure of the incision (h).

**Figure S2**

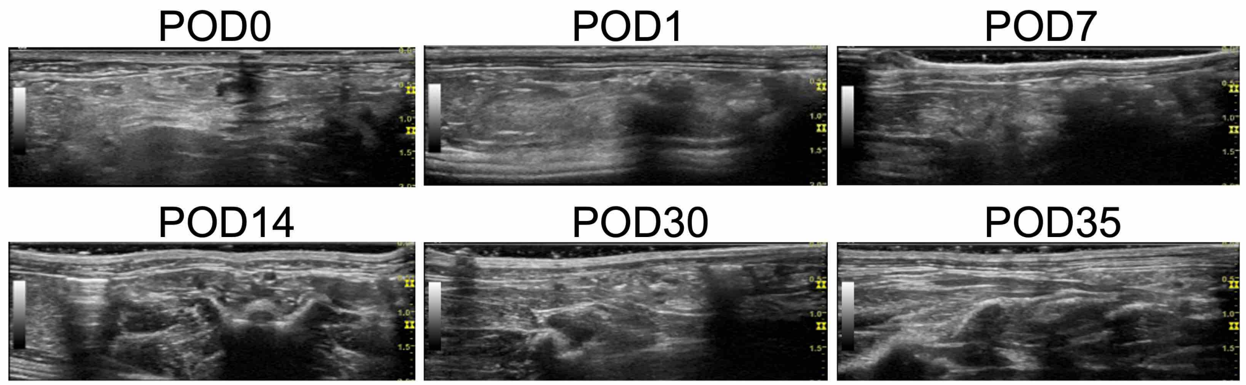


**Figure S2. Seroma-negative operated rats.** Representative longitudinal ultrasound images show no detectable fluid-filled cavity from POD0 through POD35, confirming the absence of postoperative seroma.
